# *ggtranscript*: an R package for the visualization and interpretation of transcript isoforms using *ggplot2*

**DOI:** 10.1101/2022.03.28.486050

**Authors:** Emil K. Gustavsson, David Zhang, Regina H. Reynolds, Sonia Garcia-Ruiz, Mina Ryten

## Abstract

**Motivation:** The advent of long-read sequencing technologies has increased demand for the visualisation and interpretation of transcripts. However, tools that perform such visualizations remain inflexible and lack the ability to easily identify differences between transcript structures. Here, we introduce *ggtranscript*, an R package that provides a fast and flexible method to visualize and compare transcripts. As a *ggplot2* extension, *ggtranscript* inherits the functionality and familiarity of *ggplot2* making it easy to use.

**Availability and implementation:** *ggtranscript* is available at https://github.com/dzhang32/ggtranscript, DOI: https://doi.org/10.5281/zenodo.6374061

## Introduction

Alternative splicing is a crucial post-transcriptional step through which introns are excised from messenger RNA (mRNA) precursors, and exons are spliced together to form mature mRNA isoforms. In fact, ∼95% of human genes undergo alternative splicing resulting in various forms of mature mRNA.^1^ This process is often regulated in a tissue-specific, disease-specific or developmental manner, resulting in multiple different transcripts being generated from the same gene.

It is well-recognised that standard transcriptomic assays relying on short-read RNA-sequencing are unable to comprehensively capture the full spectrum of alternative splicing.^2^ However, long-read sequencing platforms such as PacBio and Oxford Nanopore have transformed the field and enabled the discovery of new transcript isoforms that could not have been recognised by the assembly of short-reads.

Current tools to visualize transcript structures such as IGV Browser^1^, *SWAN*^2^, *wiggleplotr, Gviz*^3^, *ggsashimi*^4^ and *IsoformSwitchAnalyzeR*^5^ are often inflexible, either requiring specific data structures as input or allowing users very limited control over the outputted plot aesthetics.

Here, we introduce the R package *ggtranscript* which makes it easy to both visualize and compare transcript structures using *ggplot2*^8^. As a *ggplot2* extension, *ggtranscript* inherits a vast amount of flexibility when determining the plot aesthetics, as well as interoperability with existing *ggplot2* geoms and *ggplot2* extensions. Furthermore, the input data for *ggtranscript* matches widely used formats in genetic and transcriptomic analyses.

## Implementation

*ggtranscript* is an R package released that extends the incredibly popular tool *ggplot2* (RRID:SCR_014601 version: 3.3.5, https://cran.r-project.org/web/packages/ggplot2/index.html) for visualizing transcript structure and annotation.

As a *ggplot2* extension, the input data for *ggtranscript* is required to be a `data.frame` with columns specifying the start and end positions of each feature (e.g. exon or intron) as well as identifiers for the transcript(s) to be plotted. This data format is widely used across transcriptomic and genetic analyses and matches annotation and data structures such as the GTF/GFF3 files or `GenomicRanges` objects.

To enable the visualization of the transcript structures, *ggtranscript* introduces five new geoms (`geom_range()`, `geom_half_range()`, `geom_intron()`, `geom_junction()` and `geom_junction_label_repel()`) and several helper functions designed to facilitate the visualization of transcript structure and annotation.

`geom_range()` and `geom_intron()` enable the plotting of exons and introns, the core components of transcript annotation (**Fig 1A**). *ggtranscript* also provides the helper function `to_intron()`, which converts exon co-ordinates to the corresponding introns. Together, *ggtranscript* enables users to plot transcript structures with only exons as the required input and only a few lines of code. `geom_range()` is designed to be used for any range-based genomic annotation. For instance, when plotting protein-coding transcripts, `geom_range()` can be used to visually distinguish the coding regions from untranslated regions (**Fig 1A**).

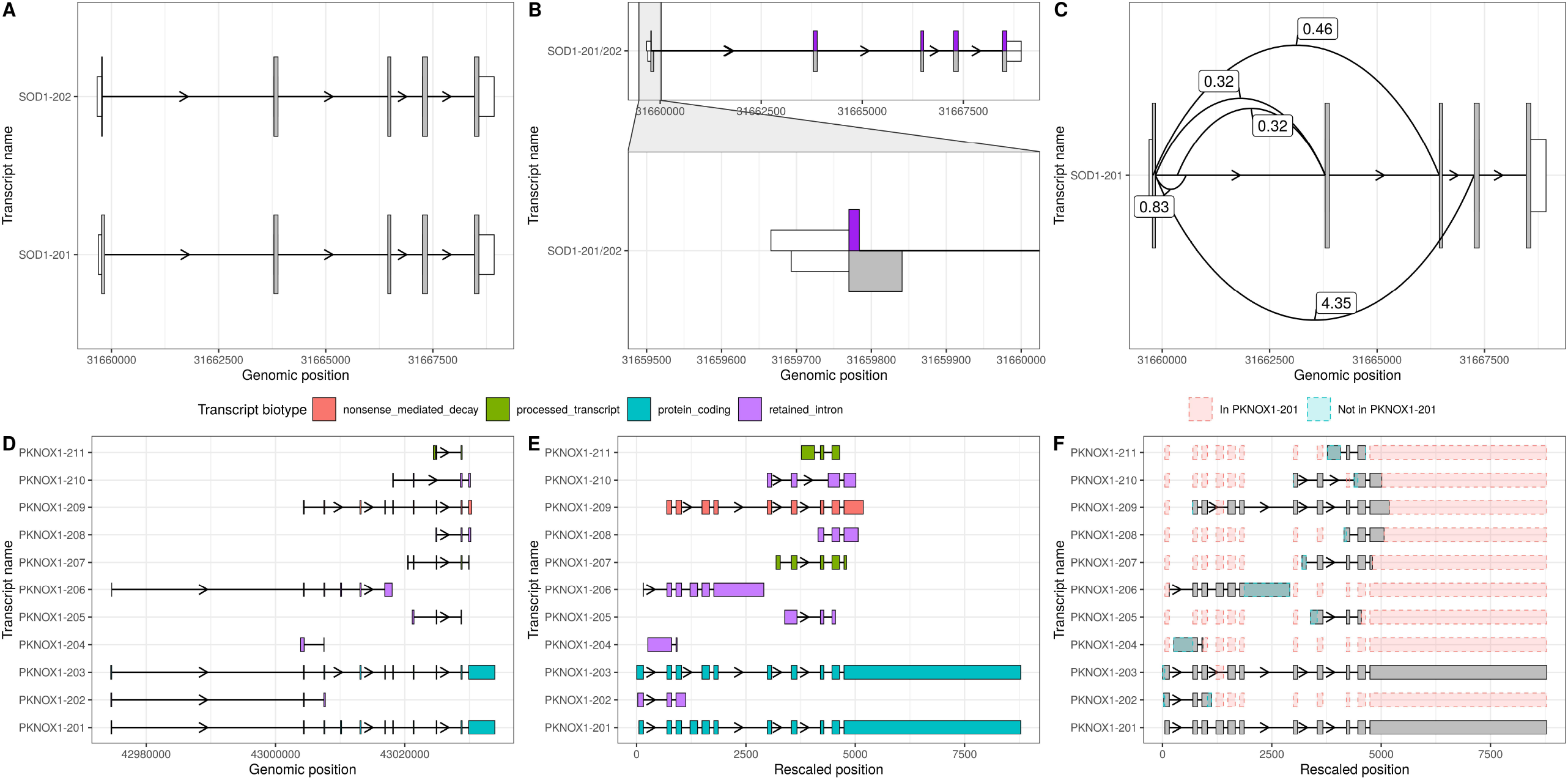
*ggtranscript* enables a fast and flexible method to visualize and compare transcript isoforms. *ggtranscript* is an *ggplot2* extension that introduces five new geoms and a set of helper functions: **A)** `geom_range()` and `geom_intron()` enable the plotting of exons and introns, the core components of transcript annotation. In addition, `geom_range()` has been used to visually distinguish coding regions (grey) from untranslated regions (white). **B)** `geom_half_range()` enables users to plot only half of a range on the top or bottom of a transcript structure; one use case of which is to visualize the differences between two transcripts (SOD201 in purple, SOD202 in grey). **C)** `geom_junction()` enables the plotting of junction curves, which can be overlaid across transcript structures to annotate them with supporting short-read RNA-sequencing data. The number represent junction usage. **D)** Longer, more complex transcripts, with small differences between exons of interest, can be more difficult to visualize. **E)** For this reason, *ggtranscript* includes a helper function `shorten_gaps()` which shortens regions that do not overlap an exon to a fixed, user-inputted width. Transcripts in D and E are coloured by their transcript biotype. **F)** In addition, the function `to_diff()` facilitates visualisation of longer transcripts by highlighting differences in comparison to a reference transcript.

`geom_half_range()` takes advantage of the vertical symmetry of transcript annotation by plotting only half of a range on the top or bottom of a transcript structure; one use case of `which` is to visualize the differences between two transcripts more clearly (**Fig 1B**). As a *ggplot2* extension, *ggtranscript* inherits the familiarity and functionality of *ggplot2*. For instance, by leveraging `ggforce::facet_zoom()` users can zoom in on regions of interest (**Fig 1B**). `geom_junction()` enables the plotting of junction curves, which can be overlaid across transcript structures to annotate them with supporting short-read RNA-sequencing data. (**Fig 1C**). `geom_junction_label_repel()` adds a label to junction curves, which can often be useful to mark junctions with a metric of their usage such as read counts (**Fig 1C**).

For longer, more complex transcripts, small differences between exons of interest can be more difficult to visualize (**Fig 1D**). For this reason, *ggtranscript* includes a helper function `shorten_gaps()` which shortens regions that do not overlap an exon to a fixed, user-inputted width. Plotting of the rescaled exons and introns enables easier comparison between transcript structures when genes are long (**Fig 1E**). In addition, the function `to_diff()` facilitates this by highlighting differences in comparison to a reference transcript (**Fig 1F**).

Together, *ggtranscript* simplifies the process of visualizing and comparing transcript structures, facilitating the exploration, analyses and interpretation of long-read sequencing and transcriptomic data.

## Conclusion

*ggtranscript* enables a fast and simplified way to visualize, explore and interpret transcript isoforms. It allows users to combine data from both long-read and short-read RNA-sequencing technologies, making systematic assessment of transcript support easier. Finally, by being a *ggplot2* extension it is highly flexible and can easily generate high-quality and publication-ready plots.

## Acknowledgements

The authors thank Siddharth Sethi and Geo Pertea for their valuable feedback and suggestions.

## Data availability

*ggtranscript* is an R package available at https://github.com/dzhang32/ggtranscript (DOI: https://doi.org/10.5281/zenodo.6374061) via an open-source MIT license. Further information and the R code of the example presented in this paper are available at https://dzhang32.github.io/ggtranscript/.

## References

1. Wang, E. T. et al. Alternative isoform regulation in human tissue transcriptomes. Nature (2008) doi:10.1038/nature07509.

2. Conesa, A. et al. A survey of best practices for RNA-seq data analysis. Genome Biol. 2016 171 17, 1–19 (2016).

3. Katz, Y. et al. Quantitative visualization of alternative exon expression from RNA-seq data. Bioinformatics 31, 2400–2402 (2015).

4. Li, Y. I. et al. Annotation-free quantification of RNA splicing using LeafCutter. Nat. Genet. 2017 501 50, 151–158 (2017).

5. Bioconductor - wiggleplotr. https://bioconductor.org/packages/release/bioc/html/wiggleplotr.html.

6. Vitting-Seerup, K., Sandelin, A. & Berger, B. IsoformSwitchAnalyzeR: analysis of changes in genome-wide patterns of alternative splicing and its functional consequences. Bioinformatics 35, 4469–4471 (2019).

7. Reese, F. & Mortazavi, A. Swan: a library for the analysis and visualization of long-read transcriptomes. Bioinformatics 37, 1322–1323 (2021).

8. Wickham, H. ggplot2: elegant graphics for data analysis. (springer, 2016).

